# Protected area establishment in Southern and Eastern Africa: Consequences for management today

**DOI:** 10.1101/2024.01.20.576490

**Authors:** Bradley Schroder, Frank Van Langevelde, Nicola-Anne Hawkins Schroder, Herbert H. T. Prins

**Author notes:** Corresponding author: Dr Bradley Schroder.

## Abstract

To understand the complexities of managing protected areas, it is important to understand the causes for their established. We summarized the motives for establishing protected areas in Southern and Eastern Africa, and the possible consequences for management of these areas today. We scrutinised documents for 48 randomly selected protected areas and investigated, (1) when and why each of the protected areas was established? (2) what the management implications were of the reasons for incorporation for these areas? and (3) how/if the original management still impacts conservation in these areas today? First, we learnt that the establishment of protected areas occurred in three periods, namely, *Period 1* when area protection was started to protect wild animals from decimation; *Period 2* during which areas considered marginal for agriculture, prone to disease or sickness and considered uninhabitable for humans were set aside; and *Period 3* when areas were proclaimed protected because of ecological or cultural importance. Second, we showed that the establishment of protected areas has ramifications for management of these areas today, which for Period 1 were remote logistics and tourism, anti-poaching difficulties, large size logistics and human wildlife conflict. Period 2 has consequences for community-land issues and intensive management, with Period 3 having intensive management to meet the objectives of these parks. Our insights have consequences for management of protected areas today, with Period 1 protected areas generally being managed on a laissez-faire approach and Period 2 and 3 protected areas being managed on a more intensive management basis.

## Introduction

“Only 50 years ago, man had to be protected from the beasts; today the beasts must somehow be protected from man” (Beard, 1988). Biodiversity is declining rapidly, caused by a variety of factors including human population growth, habitat loss, intensification of agriculture, urban sprawl, alien plant invasion and plantation forestry (Cousins et al., 2008; Huang et al., 2016; Sandbrook et al., 2019) which has the potential to threaten human life (Buitenland, 2019) but also nature. In Africa, subsistence hunting was undertaken since the Palaeolithic period, with populations of many wild animals, in Southern and Eastern Africa, being decimated especially after the commencement of a market driven economy and associated human population boom (Spinage, 1973; Waller, 1985; Carruthers, 2008; Cioc, 2009; Zulu, 2015). Early writers, missionaries and traders described Southern and Eastern Africa as the ‘nursery grounds’ for wild animals (game) and a hunter’s paradise (Elton, 1873; Wagner, 1980; Cioc, 2009). It must be recognised that most literature produced at that time in Africa were diaries written by Europeans, so we may have a distorted view point of the history of game utilisation due to the African people not documenting observations; indeed, much African history is captured in oral narration and will never be known in detail (Waller, 1985; Nash & Endfield, 2002) unless carefully analyzed (e.g., Kesteloot, 1992; Giles-Vernick, 2011). For many African peoples, game had a symbolic meaning with the most valuable animal products such as leopard skins and ivory being reserved for special use by the kings and chiefs (Zulu, 2015) and hunting itself was primarily a subsistence activity (Duffy, 2014).

During the 1830s and 1840s in Southern Africa, the “Voortrekkers” (Dutch for pioneers, also called Boers) moved away from the coastal colonies under British rule to establish independent states in the interior and flee foreign occupation. As the Voortrekkers moved inland, their lifestyles depended on and revolved around hunting and pastoralism (Plug et al., 2000; Zulu, 2015). This coupled with the so-called hunting expeditions by European “sportsmen” and the ivory trade, led to the largest slaughter of wildlife in the mid-nineteenth century (Spinage, 1973; Murombedzi, 2003; Carruthers, 2008; Cioc, 2009; Zulu, 2015). This was by no means only a problem associated with Africa; for example, vast herds of bison were decimated in North America due to hunting and the establishment of human settlements (Muir-Leresche & Nelson, 2000; Child et al., 2012).

In the first half of the twentieth century, it was considered common practice in Southern and Eastern Africa, that game should be exterminated to make way for domestic animals such as cattle, sheep and other domestic species (Murombedzi, 2003; Carruthers, 2008; Cioc, 2009). In 1903, for example, the government of Kenya introduced a scheme whereby settlers were paid to kill certain species of wild animals, especially carnivores (Murombedzi, 2003). The various agricultural and game departments acted accordingly and decimated game throughout Southern and Eastern Africa (Carruthers, 2008). For example, in Rhodesia (now Zimbabwe) and Zambia, it became an official government policy to eliminate wildlife wherever it was perceived to threaten or compete with agriculture in any way or form.

Not only competition for resources and predation of livestock resulted in a decline in wild animal populations, disease control implemented for diseases such as bovine pleuropneumonia (1850), rinderpest (1896) and tsetse fly, also lead to game decimation (Mutwira, 1989; Cioc, 2009). The idea that wild animals were the primary vectors of disease, almost brought wildlife to extinction in Southern and Eastern Africa. In the Zambezi and Savé valleys between 1919 and 1958, for example, over 660,000 head of game were shot as part of disease control (Child & Riney, 1987), and between 1919 to the 1980s, thousands of wild animals including rhinoceroses were shot in the name of tsetse fly control (Taylor & Martin, 1987; Prins, 1996). Moreover, during the 1980s, large areas were fenced to control foot-and-mouth disease in a number of Southern African countries (Child et al., 2012), which lead to animals dying from starvation and lack of water (especially migratory animals).

It was becoming obvious by the late eighteenth century that the continued decimation of wild animals could not be sustained, or extinction of the majority of species would be imminent (Carruthers, 1988; Gibson & Marks, 1995). In pre-colonial Botswana, for example, the reduction of animal populations led to a monopolization of access to wildlife; the Tswana chiefs were given control over traditional, foreign commercial and sport hunting (Hitchcock, 2001; Neumann, 2002; Murombedzi, 2003; Pangeti & Manyanga, n.d.), which may have led to a form of protection. In the late 1800s a number of people including conservationists, sports hunters and military personnel decided that wild animals needed protection to ensure their survival and for sustainable utilisation in the future. Pre-colonial conservation in South Africa was devised by means of several strategies for conserving nature while at the same time guaranteeing access to it (Murombedzi, 2003). Examples of these are the Sabi Game Reserve which was established to protect animals for hunting (this was a decision made by the Volksraad in 1898), Kruger National Park started through political agendas, and the Bontebok National Park started to protect the Bontebok species. The cutting edge of conservation at that time lay in South Africa, with the establishment of the continent’s first game reserve in 1895, Hluhluwe-Imfolozi, in Zululand. This was followed by the first protected area in 1898, the Sabi Game Reserve, which in 1926 became the country’s first National Park and was renamed, the Kruger National Park (Rangarajan, 2003).

In the beginning of the twentieth century, there were three policy decisions taken in South Africa which would have long-lasting implications for conservation. Firstly, promotion of the formation of state protected areas; secondly, restriction of the commercial use of wildlife, rendering wildlife valueless except for low-value subsistence uses; and thirdly, centralization of ownership of wildlife in the state, disenfranchising landholders and taking upon themselves the burden of protecting wildlife from people (anti-poaching) and protecting people from wildlife (problem animal control) (Heijnsbergen, 1997; Child et al., 2012). While the origins of European interest in Southern and Eastern African wildlife conservation can be traced to the activities of aristocratic hunter-naturalists, late colonial Africa became the proving ground for political and technical strategies of conservation organisations that are today key parts of global environmental management (Neumann, 2002). This was supported by European countries in 1900, at the conference of the “Preservation of Wild Animals, Birds and Fish in Africa”, held in London (Neumann, 2002; Coic, 2009) and in 1933 at the conference of the “Protection of the Fauna and Flora of Africa” (Neumann, 2002). The conference only included certain African countries under European control and the majority of people involved in the conferences came from British or European aristocracy and all big game hunters. The conference primarily focussed on what they called the “indiscriminate slaughter” of wildlife by Africans and their recommendations were to outlaw most customary African hunting practices. It only protected a handful of animals, declared subsistence hunting as poaching and only allowed hunting of wild animals with a permit (Mutwira, 1989; Grobler, 1996; Cioc, 2009). Yet, many efforts failed and even the leading politician of the Union of South Africa (General Jan Smuts) was unable to sway land owners, miners and the civil administration into the creation of a large protected area centred on what is now the Mapungubwe National Park in the 1940s (Carruthers, 1992).

There were many changes to conservation policies over the coming century but it was in the early 2000s, that the South African Government noticed numerous flaws in the legal framework for the identification, declaration and management of the majority of protected areas, which led in 2003 to the overhaul of the country’s national conservation regime with the establishment of the National Environmental Management: Protected Areas Act 2003 (NEM:PAA) (Paterson, 2009).

It became clear over the years that nature conservation and the protection of natural areas were initiated based on different motives, which may have consequences for the management of the land and biodiversity which is protected today (Renwick & Schellhorn, 2016). In this paper, we analyse the establishment of protected areas in Southern and Eastern Africa to understand the motives for establishing the protected areas and the possible consequences for management of these areas today.

### The protection of natural resources during the colonial era

During the colonial era, in Southern and Eastern Africa there were similar motives for the establishment of protected areas. To illustrate this, we describe the developments in a number of countries, i.e., Sudan (in what is now South Sudan), Rhodesia (now Zambia and Zimbabwe) and Kenya. In the Sudan, for example, the decimation of wildlife started with colonial sport hunting parties such as those undertaken by Roosevelt in 1909, which brought the concept of the ‘safari’ to international attention (Duffy, 2014). In more recent times, the bush meat trade and especially the years of warfare in which wild animals were used to feed soldiers, had a large toll on wildlife. Rhodesia based its game legislation on Roman Dutch law, which established that wild animals were “*res nullius*”, which is “a thing or things which can belong to no one” (Oxford Dictionary, 2017). This resulted in wild animals being seen as “Royal Game”, to be held by the king (of Britain), in his “sovereign capacity for the people” (we will not delve into the justification of that: see for instance Strang (1996) or Culhane (1998) but from the colonial power’s point of view it was based on its concept of *lus publicum* (public law) (Marston, 1996), even though that was perhaps merely a fictitious concept to justify resource grabbing by right of conquest (Kirby & Kirby, 1931)). In 1889, the policy of game preservation was written into the Charter of Incorporation, which signalled the beginning of game legislation in Rhodesia. The ordinance declared the preservation of certain wild animals, specifically elephant (*Loxodonta africana*) and that no interference with hunting rights bestowed on native chief’s or peoples could be undertaken unless to enforce closed seasons (Charter of the British South Africa Company, 1889). The Game Preservation Ordinance, No 6 of 1899, was the first official law to protect wildlife and declared species such as elephant, eland (*Taurotragus oryx*), giraffe (Gi*raffa camelopardalis*), gemsbok (*Oryx gazella*), hippopotamus (*Hippopotamus amphibius*), kudu (*Tragelaphus strepsiceros*), ostrich (*Struthio camelus*), square-lipped rhinoceros (*Ceratotherium simum*), hook-lipped rhinoceros (*Diceros bicornis*) and zebra (*Equus burchellii*) as “Royal Game” and prohibited their killing, hunting or capture, unless permission was granted by the administration (Mutwira, 1989). The Game and Fish Preservation Act in Rhodesia (No. 35 of 1929) allowed for the establishment of game reserves, and in 1929 the Matopo National Park was established as Rhodesia’s first national park. This was intended to preserve all species of flora and fauna occurring in the designated area, which would be protected from hunting and used for scientific, educational and aesthetic purposes (Murombedzi, 2003). The 1930s saw a vast increase in game reserves, the appointment of wardens and the legislation of private game reserves (Mutwira, 1989). The Department of National Parks was established in 1949, which led to the single most important change in Rhodesia’s conservation history, the promulgation of the Parks and Wildlife Act in 1975, which conferred privileges on the owners and occupiers of alienated land as custodians of wildlife.

In 1899 and 1900, respectively, the Kenyan government established the Southern and Northern Game Reserves, covering nearly 70,000 km^2^, and the establishment of the Game Department in 1907 to protect reserves and enforce game laws. As with most other game departments of the time, this was hampered by a lack of money, insufficient and unqualified staff. The Kenya Game Department established a “vermin policy” which eradicated all species seen to be a threat to humans or agriculture, both inside and outside of protected areas. This included but was not limited to species such as lion (*Panthera leo)*, leopard (*Panthera pardus*), hyaena (*Crocuta crocuta*) and wild dog (*Lycaon pictus)*, with devastating effects on these species (Carruthers, 1988; Mutwira, 1989; Waithaka, 2012). In 1932, the British government decided on the establishment of “National Parks and reserves where hunting, killing or capturing of fauna, and the collection or destruction of flora would be limited or prohibited”, which resulted in the establishment of the 117 km^2^ Nairobi National Park in 1946.

Wildlife formed a focus for the divergent interests of the black Africans, Afrikaners and British in South Africa (Carruthers, 1988). As rural African populations grew, local subsistence hunting was deemed to threaten wildlife and this was treated as poaching (Mutwira, 1989). Poachers were prosecuted by the colonial authority of the time, which embedded military approaches and values in modern conservation practices, this has become known as “green militarization” (Mutwira, 1989; Dahlberg et al., 2010; Duffy, 2014; Hitchcock, 2019). The local populations saw this traditional style of livelihood as a right and by removing their hunting rights it established a sense of injustice and negativity towards protected areas, wildlife and the state (Gibson & Marks, 1995; Child et al., 2012). At the twilight of colonial Africa, many leading conservationists met at the Arusha Conference on the ‘Conservation of Nature and Natural Resources in Modern African States’, realizing that a new post-colonial narrative was needed to conserve wildlife (Watterson, 1961) and that wildlife needed to become an economic asset according to the emerging ‘use it or lose it’ philosophy. This was formalized at the Conference of Parties of the Convention on Biological Diversity (1998) by all Member States accepting the Principles of Sustainable Use.

During the period from 1970 to 2005, a number of national parks and wildlife reserves across Africa lost up to 60% of their wild animals. Researchers studied animal population changes on 78 protected areas across Africa and found that West Africa had lost up to 85% of its wildlife in 35 years, and East Africa nearly 50% (Ottichilo et al., 2000; Ottichilo et al., 2001; Vidal, 2011). It was only found that wildlife numbers increased in Southern Africa (Child et al., 2012). There is a large amount of speculation as to why there has been a decline in wildlife numbers across the continent and with limited scientific proof as to the exact causes. It is certain that politics, historically poor management of conservation areas, lack of funding, lack of qualified personnel, lack of political support, human activities encroaching on wildlife areas, wars, bush meat trade, veterinary interventions and poaching are all severe causes of the decline (Mbaiwa & Mbaiwa, 2006; Cioc, 2009; Vidal, 2011). Growing human populations, unsustainable hunting, high densities of livestock, and habitat loss have devastating consequences for large herbivore species, their ecosystems, and the services they provide (Ottichilo et al., 2001; Ripple et al., 2015).

### Different periods of the establishment of protected areas in Southern and Eastern Africa

Given the various timelines and explanations for the establishment of protected areas as given in this overview, we defined the establishment of protected areas in Southern and Eastern Africa in three “periods”. It is known that not all countries have identical timelines in history, thus we distinguish “periods” which are not necessarily in synchrony in the different Southern and Eastern African countries but which we believe all these countries undergo.

We identify *Period 1* as the establishment of protected areas that were initially started to protect wild animals from decimation for utilisation and enjoyment in the future (primarily by sports hunters) (Carruthers, 1988; Murombedzi, 2003). An example of Period 1 is the Sabi Game Reserve in South Africa, established in 1898, later to become known as the Kruger National Park (1926). With the establishment of the continent’s first national park, the Kruger National Park, conservation paradigms throughout the continent had begun to change. In the Kruger Park, sports hunting was forbidden in an attempt to protect the last remaining wild animals. Such an act of creating a park was not merely a gesture to save nature, but to redefine who owned it (Rangarajan, 2003). The creation of many protected areas around the world often resulted in the alienation of indigenous populations from their land and resources (Kepe et al., 2005; Hitchcock, 2019).

*Period 2* is identified as areas that were initially considered marginal for agriculture, prone to disease or sickness, considered uninhabitable for humans, with a struggling yet established wildlife populations (Department of Environmental Affairs and Tourism, 1997; Paterson, 2009). Due to the marginal land not being sustainable for agriculture and/or farming, it was used for hunting and tourism in order to secure an income. An example of this type of protected area is the St Lucia Game Reserve in Zululand (South Africa), established in 1895. The locations of these types of protected areas were pre-determined by the presence of Tsetse fly and malaria, hence unsuitable for livestock, or by the fact that their agricultural potential was poor (Department of Environmental Affairs and Tourism, 1997).

*Period 3* only commenced much later, with the promulgation of the first National Parks Act in South Africa, when protected areas were proclaimed for ecological or cultural importance, such as the protection of a cultural site, a specific threatened species (animals and vegetation) or a water source (Paterson, 2009). Examples are the Bontebok and Kalahari National Parks in South Africa in 1931 and Gonarezhou National Park (Zimbabwe), in 1975.

## Methods

To better understand the three periods of establishment of protected areas in Southern and Eastern Africa, 48 protected areas from various countries (the countries which form part of the study are Botswana, Kenya, Namibia, South Africa, Tanzania, Uganda, Zambia and Zimbabwe) were randomly selected to compare certain criteria and principles in an attempt to understand the following three main questions, (1) when and why each of the protected areas was established? (2) what the management implications were of the reasons for incorporation for these areas? and (3) how/if the original management still impacts conservation in these areas today?

We only selected protected areas that are situated in the savanna biome. Savannas represent one of the largest biomes of the world, comprising roughly 20% of the earth’s land area (Shorrocks & Bates, 2007; Huntley & Walker, 2012). Savannas in Africa occupy almost 50% of the land area of the continent and support not only a large portion of its humans, and livestock, but some of the highest densities and diversity of wild herbivores and carnivores in the world (Scholes & Archer, 1997; Shorrocks & Bates, 2007; Sankaran & Anderson, 2009). This study has no relevance to rain forests or desert areas as we are looking at protected areas in savannas that are under a higher threat from human populations and their requirements for arable land. The 48 protected areas were randomly selected using Google search (key words – Africa, Southern, Eastern, Conservation, Protected, Areas). Both the English and Afrikaans languages were used in the literature research criteria. 689 articles, 25 books and 31 management plans of these 48 protected areas were traced and read. The management plans were either obtained from the protected area directly or from search engines. Lusophone and Francophone countries were not included in the search criteria to ensure that protected areas for only Southern (excl. Mozambique and Angola) and Eastern Africa (excl. Rwanda and Burundi) were obtained. Based on the historic overview, we assumed that protected areas established in Period 1 would be established on an ad-hoc basis, primarily to protect animals for later use and that they would generally be larger than protected areas in the other two periods. The protected areas in this period would have been established on land previously used for hunting. We assumed that protected areas in Period 2 would be established on marginal lands, previously used for agriculture and/or domestic animal farming. For those protected areas we expected certain forms of manipulation as nutrient rich vegetation for large mammalian herbivores is not prolific on marginal lands. Period 3 protected areas would have been established on conservation and/or biodiversity principles with previous land use being cultural uses of one form or another.

The following sub-questions were addressed for each protected area to obtain a better understanding of the three main questions.

- What were the primary motives for the establishment of the protected area? (Question 1)
- What was the previous land use prior to establishment of the protected area? (Questions 2 and 3)
- What are the consequences for present day management based on the site selection of the protected area? (Question 3)
- What is the main current attraction of the protected area? (Questions 2 and 3)
- Has the management of the protected area used manipulation of animals and/or vegetation as a management tool? (Questions 2 and 3)

The primary motives for establishing the protected areas (Table 1) varied, with the majority being established for tourism and the protection of animal species. Conservation and biodiversity only ranked third. Parks established with protection of animal species, hunting, scenery, geography and game breeding as the primary motives for establishing protected areas were assigned to Period 1 (Table 1). The protected areas that were established more for tourism, hunting, education, joint conservation initiatives and to protect lions were assigned to Period 2. Period 3 protected areas were established for conservation and/or biodiversity, community and/or culture, education and for the protection of elephants and rhinoceroses.

**Table 1.** Motives as published in reports as the rationale for the establishment of the 48 protected areas. For the 48 protected areas we investigated, multiple motives for their establishment were reported, yielding a total of 83 motives. This has been tabulated in the second column “primary motive for establishment”. The ratio per phase indicates whether the primary motive for establishment was “preferred” (>1) or “avoided” (<1). The three identified periods are represented as P1, P2 and P3

| Primary motives (n = 13) for the establishment of the protected areas | Frequency with which a motive is given for the establishment of a protected area | The protected areas using the individual primary motive for establishment for each period and the relevant percentage | Ratio per period indicating whether the reason was “preferred” (>1) or “avoided” (<1) |
| --- | --- | --- | --- |
| Tourism | 25 | P1 : 16%<br>P2 : 76%<br>P3 : 8% | P1 : 0.55<br>P2 : 1.52<br>P3 : 0.38 |
| Protect animal species | 24 | P1 : 38%<br>P2 : 46%<br>P3 : 17% | P1 : 1.29<br>P2 : 0.92<br>P3 : 0.79 |
| Conservation & Biodiversity | 14 | P1 : 29%<br>P2 : 29%<br>P3 : 43% | P1 : 0.99<br>P2 : 0.57<br>P3 : 2.04 |
| Community & Culture | 3 | P2 : 33%<br>P3 : 67% | P2 : 0.67<br>P3 : 3.17 |
| Hunting | 3 | P1 : 33%<br>P2 : 67% | P1 : 1.15<br>P2 : 1.33 |
| Scenery | 3 | P1 : 100% | P1 : 3.45 |
| Education | 2 | P2 : 50%<br>P3 : 50% | P2 : 1.00<br>P3 : 2.38 |
| Geography | 2 | P1 : 100% | P1 : 3.45 |
| Joint conservation initiative | 2 | P2 : 100% | P2 : 2.00 |
| Protect elephants | 2 | P3 : 100% | P3 : 4.76 |
| Game breeding | 1 | P1 : 100% | P1 : 3.45 |
| Protect lions | 1 | P2 : 100% | P2 : 2.00 |
| Protect rhinoceros | 1 | P3 : 100% | P3 : 4.76 |

The previous land use for the majority of protected areas fell into four main categories, colonial hunting, ranching, agriculture and indigenous hunter-gatherers (Table 2). Period 1 protected areas were used for colonial hunting, indigenous hunter-gatherers, fishing and iron smelting. The Period 2 protected areas were used for cattle ranching (and other ranching) and rhinoceros conservation, while Period 3 protected areas were used for colonial hunting, agriculture (mixed crops), indigenous hunter-gatherers, fishing, Iron smelting and mining.

**Table 2.** Previous land uses of the 48 protected areas. For the 48 protected areas we investigated, multiple previous land uses were established, yielding a total of 105 uses. This has been tabulated in the second column “frequency of previous land use”. The ratio per period indicates whether the previous land use was “preferred” (>1) or “avoided” (<1). The three identified periods are represented as P1, P2 and P3

| Previous land use (n = 8) for the 48 protected areas | Frequency of previous land use in the 48 protected areas | The protected areas using the individual previous land use for each period and the relevant percentage | Ratio per period indicating whether the previous land use was “preferred” (>1) or “avoided” (<1) |
| --- | --- | --- | --- |
| Colonial hunting | 32 | P1 : 31%<br>P2 : 47%<br>P3 : 22% | P1 : 1.08<br>P2 : 0.94<br>P3 : 1.04 |
| Cattle ranching (and other ranching) | 29 | P1 : 10%<br>P2 : 69%<br>P3 : 21% | P1 : 0.36<br>P2 : 1.38<br>P3 : 0.99 |
| Agriculture (mixed crops) | 19 | P1 : 26%<br>P2 : 47%<br>P3 : 26% | P1 : 0.91<br>P2 : 0.95<br>P3 : 1.25 |
| Indigenous hunter-gatherers | 16 | P1 : 44%<br>P2 : 31%<br>P3 : 25% | P1 : 1.51<br>P2 : 0.63<br>P3 : 1.19 |
| Fishing | 3 | P1 : 67%<br>P3 : 33% | P1 : 2.30<br>P3 : 1.59 |
| Iron smelting | 3 | P1 : 33%<br>P2 : 33%<br>P3 : 33% | P1 : 1.15<br>P2 : 0.67<br>P3 : 1.59 |
| Mining | 2 | P3 : 100% | P3 : 4.76 |
| Rhinoceros conservation | 1 | P2 : 100% | P2 : 2.00 |

For each sub-question we found categories that were compared among the three periods. We counted the number of parks per period and per category to calculate a ratio per period and to indicate whether this category was more frequently mentioned than expected based on how many parks were assigned to each period:

a = Number of parks in a period / Total number of parks
b = Number of parks in a period that mention this category / Total number of parks that mention this category per period
Ratio = b / a

When the ratio is > 1, then the category is mentioned more frequently than expected (“preferred”), whereas the category is mentioned less frequently than expected the ratio is < 1 (“avoided”). To address the sub-questions, we only looked at ratios >1 as these ratios highlight the consequences for management of the protected areas today. We considered ratios between 1 – 1.1 as only slightly deviating from no “preference” (the baseline), ratios between 1.1 – 1.2 as having a moderate deviation and ratios >1.2 as having a large deviation from no “preference”.

## Results

We found that protected areas established during Period 1 on average (1844 x 10² km²) are considerably larger than both Period 2 (9 x 10² km²) and Period 3 (12 x 10² km²). This is primarily due to the earlier protected areas in Period 1 being established in remote areas with minimal human inhabitants at the time while in the latter years, protected areas were established for specific motives and often close to human settlements constraining the size of the protected area (see Supporting documentation A for the information pertaining to each of the 48 protected areas). Twenty-nine percent of the 48 protected areas included in this study were established in Period 1 as they were primarily established (by sports hunters) to protect wild animals from decimation, for utilisation and enjoyment in the future. The majority of the 48 protected areas in our overview (50%) were established in Period 2, and were areas considered marginal for agriculture, prone to disease or sickness, considered uninhabitable for humans, or with a struggling yet established wildlife populations. Period 3, which only became a reality after 1926 with the establishment of the National Parks Act in South Africa, were represented by 21% of the protected areas. The protected areas proclaimed in Period 3 were established because of ecological or cultural importance.

The consequences of site selection of the protected areas have varying ramifications for conservation management today. Table 3 shows current management constraints due to the original site selection with the ratios clearly illustrating which of the consequences are frequently mentioned per period.

**Table 3.** Consequences of site selection of the 48 protected area for nowadays management. For the 48 protected areas we investigated, multiple consequences of site selection were established, yielding 195 consequences. This has been tabulated in the second column “frequency of consequences of site selection”. The ratio per period indicates whether the consequence of site selection was “preferred” (>1) or “avoided” (<1). The three identified periods are represented as P1, P2 and P3

| Consequences of site selection (n = 11) for the 48 protected areas | Frequency of consequences of site selection for the 48 protected areas | The protected areas using the individual consequences of site selection for each period and the relevant percentage | Ratio per period indicating whether the consequences of site selection was “preferred” (>1) or “avoided” (<1) |
| --- | --- | --- | --- |
| Veterinary restrictions | 48 | P1 : 29%<br>P2 : 52%<br>P3 : 19% | P1 : 1.01<br>P2 : 1.04<br>P3 : 0.89 |
| Community-land issues | 23 | P1 : 26%<br>P2 : 61%<br>P3 : 13% | P1 : 0.90<br>P2 : 1.22<br>P3 : 0.62 |
| Remote logistics (hard to access) | 20 | P1 : 65%<br>P2 : 25%<br>P3 : 10% | P1 : 2.24<br>P2 : 0.50<br>P3 : 0.48 |
| Remote tourism (hard to access) | 20 | P1 : 65%<br>P2 : 25%<br>P3 : 10% | P1 : 2.24<br>P2 : 0.50<br>P3 : 0.48 |
| Intensive management | 20 | P1 : 5%<br>P2 : 65%<br>P3 : 30% | P1 : 0.17<br>P2 : 1.30<br>P3 : 1.43 |
| Anti-poaching difficulties | 19 | P1 : 74%<br>P2 : 16%<br>P3 : 11% | P1 : 2.54<br>P2 : 0.32<br>P3 : 0.50 |
| Large size logistics | 16 | P1 : 69%<br>P2 : 13%<br>P3 : 19% | P1 : 2.37<br>P2 : 0.25<br>P3 : 0.89 |
| Human animal conflict | 12 | P1 : 67%<br>P2 : 17%<br>P3 : 17% | P1 : 2.30<br>P2 : 0.33<br>P3 : 0.79 |
| Climatic issues (drought/flood) | 7 | P1 : 86%<br>P3 : 14% | P1 : 2.96<br>P3 : 0.68 |
| Military incursions | 5 | P1 : 60%<br>P2 : 20%<br>P3 : 20% | P1 : 2.07<br>P2 : 0.40<br>P3 : 0.95 |
| Poor management | 5 | P1 : 40%<br>P2 : 40%<br>P3 : 20% | P1 : 1.38<br>P2 : 0.80<br>P3 : 0.95 |

Because Period 1 includes protected areas established to protect species for later use, and due to their very large sizes, the main consequences for management today is the practise of management through the laissez-faire approach (non-interventionist approach, neither interference nor manipulations) coupled with remote logistics and tourism, anti-poaching difficulties, large size logistics, human animal conflict, climatic issues, military incursions and poor management.

Because Period 2 includes protected areas that were established on marginal or cultural land, the main consequences for management today are community-land issues and intensive management (including but not limited to bush clearing and control, alien plant control, supplementary feeding, lick blocks or vegetation restoration).

Because Period 3 protected areas were established on conservation and/or biodiversity principles, the main consequence for management today is intensive management (again including but not limited to bush clearing and control, alien plant control, supplementary feeding, lick blocks or vegetation restoration).

The current main attractions for each of these protected areas has an influence on the management goals and objectives of these protected areas today. Table 4 shows current main attractions for the 48 protected areas with the ratios clearly illustrating which of the consequences are mentioned frequently per period. It is evident that the current main attraction for the majority of areas is the iconic Big 5 (81%) (viz. lion, leopard, elephant, black rhinoceros – *Diceros bicornis* and African buffalo – *Syncerus caffer*). The nine protected areas, which did not have the Big 5, all had the Big 4 (Big 5 less the rhinoceros).

**Table 4.**
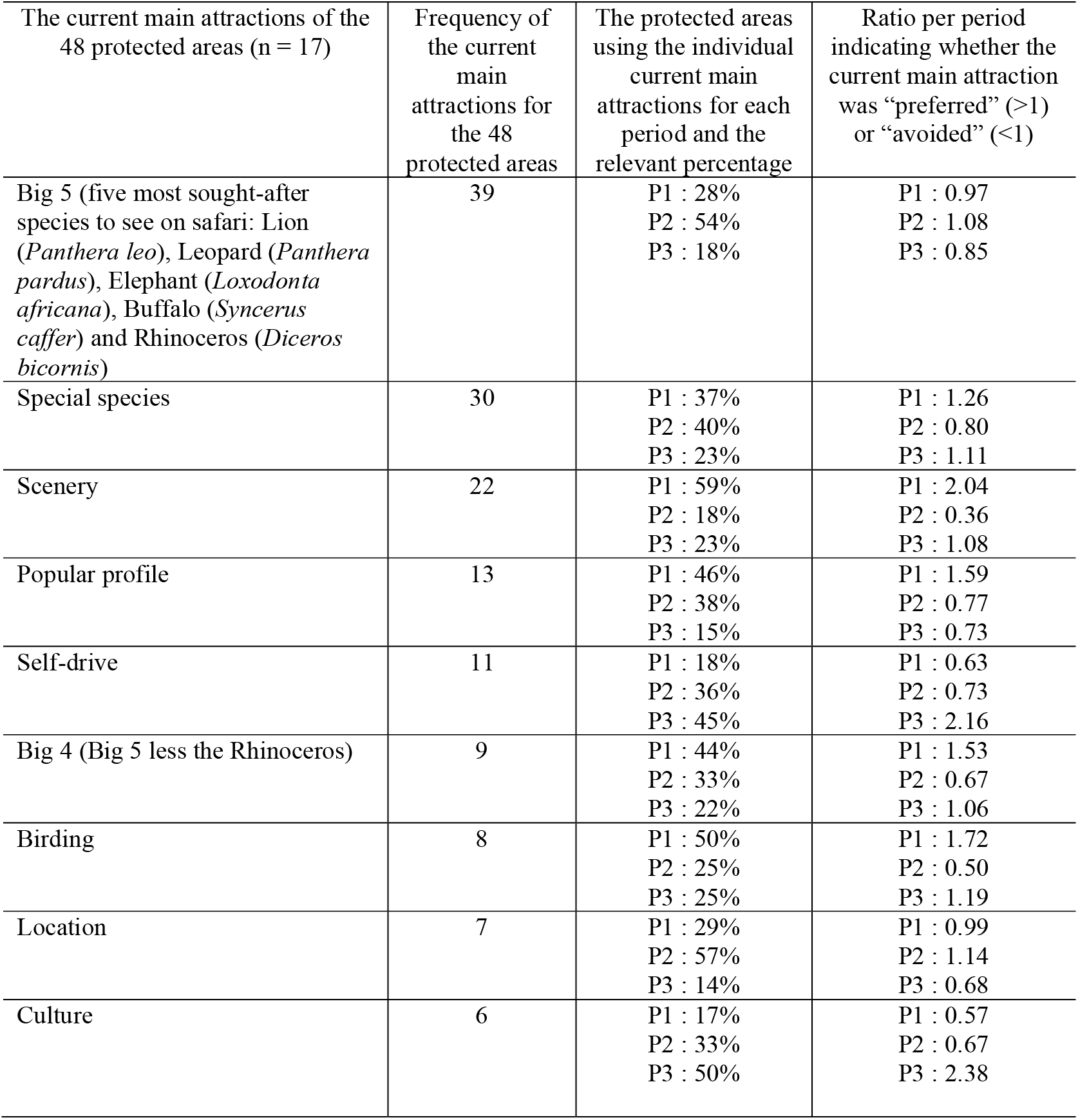
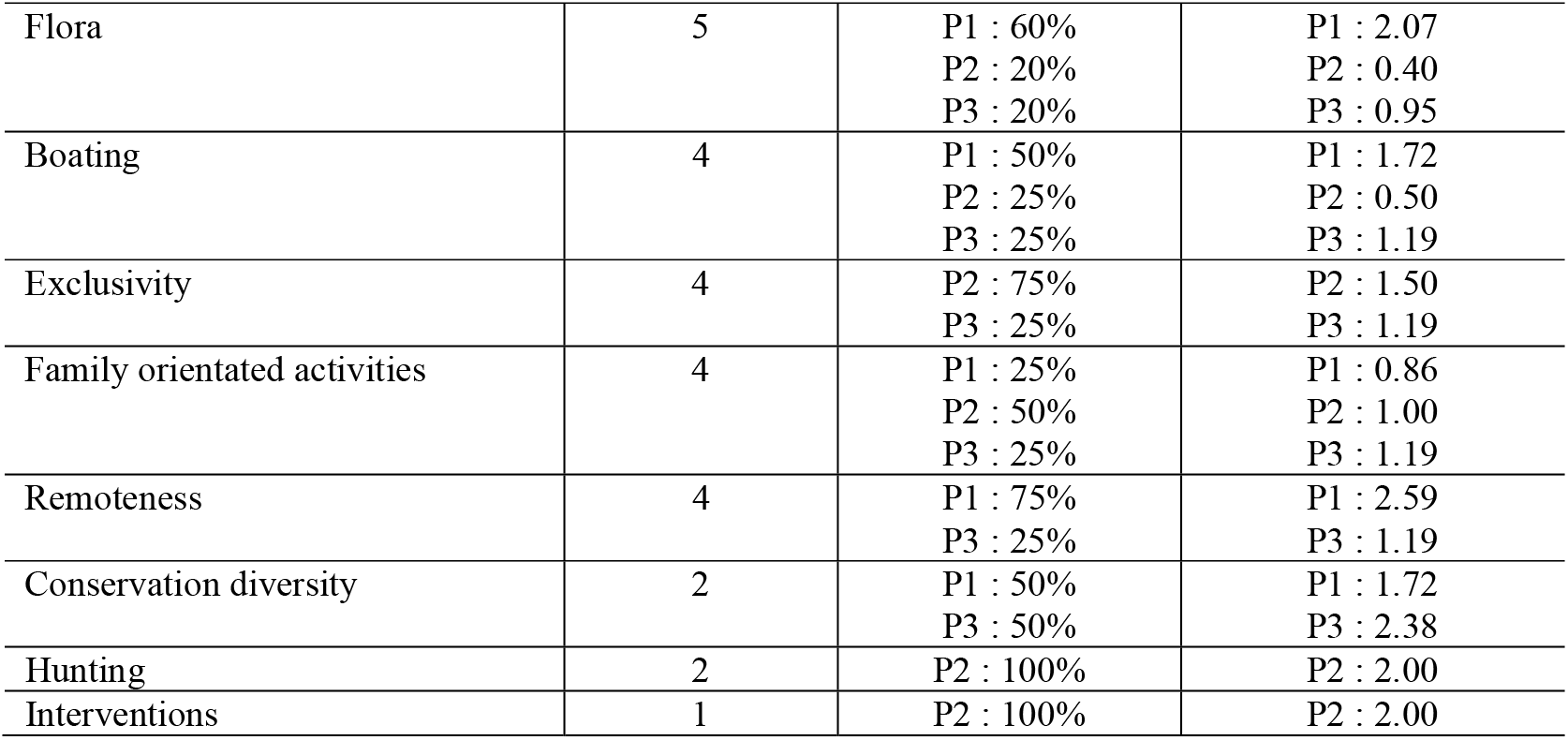
The current main attractions of the 48 protected areas. For the 48 protected areas we investigated, multiple current main attractions were established, yielding 171 attractions. This has been tabulated in the second column “frequency of the current main attractions”. The ratio per period indicates whether the current main attraction was “preferred” (>1) or “avoided” (<1). The three identified periods are represented as P1, P2 and P3

Because Period 1 includes protected areas established to protect species for later use, the main attractions are special species, scenery, popular profile, Big 4, birding, flora, boating, remoteness and conservation diversity.

Because Period 2 includes protected areas that were established on marginal or cultural land for hunting and tourism, the main attractions were Big 5, location, exclusivity, family orientated activities, hunting and interventions.

Because Period 3 protected areas were established on conservation and/or biodiversity principles, the main attractions are special species, scenery, self-drive, Big 4, birding, culture, boating, exclusivity, family orientated activities, remoteness and conservation diversity.

Of the 48 protected areas, 37 of the protected areas partook in the manipulation of animal populations or vegetation (i.e., active management) and 2 undertook minimal manipulation. We were unable to establish if the remaining 9 protected areas partook in any form of manipulation (Appendix A). The ratios illustrate that Periods 2 and 3 were identified as using manipulations as an often-mentioned reason as part of their protected areas management.

## Discussion

In this paper we discerned the motives for establishing protected areas in Southern and Eastern Africa, and the possible consequences for management of these areas today.

Based on our historical overview of the establishment of protected areas in Southern and Eastern Africa, we distinguished three periods in which these protected areas were established, the primary motives for establishing the protected areas, what the previous land uses were, the consequences of the site establishment for management today, the current attractions and whether or not they participated in animal population or vegetation manipulation.

Period 1 protected areas were primarily established to protect species for later use. Prior to being established as a protected area, the previous land uses included colonial hunting and indigenous hunter-gatherers (including fishing) and iron smelting. According to Dahlberg et al. (2010) protected areas were proclaimed primarily to protect wildlife and as a refuge from ills of civilisation and for the recreation of the human spirit. The consequences of establishing these protected areas in the locations resulted in the areas being more complicated to manage than either Period 2 or 3 for current management, from a logistics perspective based on their larger size and remoteness of the protected areas. This includes but is not limited to fence management, anti-poaching, supply chain, tourism access and poor management. These consequences are recognised for the majority of large protected areas in remote places (Worboys & Trzyna, 2015). All 14 protected areas identified in Period 1 are managed by the relevant National Parks Board Authorities of these countries and manipulation (active management) of the populations of animals and/or the vegetation is not viewed as a preferred strategy when it comes to management planning. Due to the size of the areas, animals and vegetation are generally managed on a laissez-faire approach (non-interventionist approach, neither interference nor manipulations). This is in contrast to earlier stages when in remote areas within the protected areas predator control was the norm for a long-time (Stevenson-Hamilton, 1938). One should also note that in many of the protected areas where laissez-faire is advocated, the construction of artificial water points is a very intense form of management and has been executed for decades (Hilbers et al., 2015) and burning is considered acceptable even if the consequences are largely ignored (Groen et al., 2008; Klop & Prins, 2008; Masocha et al., 2011). The current main attractions for these protected areas include conservation biodiversity which includes special species, scenery, birding and flora, coupled with these areas being attractive for their popular profiles and remoteness. Due to these protected areas being owned by government, the areas are not as intensively driven by economic survival as there is state funding to ensure the longevity of these protected areas.

Period 2 protected areas were primarily established for tourism and hunting. Prior to being established as a protected area, the previous land uses were dominated by cattle ranching (and other ranching). Enghoff (1990) states that wildlife as a form of land use must be considered as one of the major forms of alternative use for semi-arid pastoral land. The consequences of establishing these protected areas in their locations resulted in the areas being less complicated to manage from the perspective of size and location but due to the marginal land the requirement for intensive management of both animal populations and vegetation increased, with higher rates of issues pertaining to local communities. Seventy-one percent of the protected areas identified in Period 2 are privately owned and managed. Manipulation of the animal populations and vegetation in one way or another is often mentioned when it comes to management planning, primarily due to the marginal land. According to Ajathi & Krumme (2002), anthropogenic disturbances in natural ecosystems are of a foreign character with vast ecological consequences in degradation. The current main attraction for these protected areas is tourism (Big 5). Due to these protected areas being owned and managed by private entities they are driven by tourism as the primary form of economic survival to ensure the longevity of these protected areas, resulting in the management today focusing their efforts on the tourism portfolio. James et al. (1999), posit that effective management of protected areas depends greatly upon adequacies of available resources and that smaller protected areas require great budgetary and staffing inputs.

Period 3 protected areas were established primarily on the principles of conservation and/or biodiversity and community and/or culture. The past century has seen a vast increase in the establishment of protected areas, to protect biodiversity and refuges for particular species to entire ecosystems (Rao et al., 2009). Prior to being established as a protected area, the previous land uses included colonial hunting and agriculture (mixed). Due to all the protected areas in Period 3 being established after 1926, with the establishment of the National Parks Act in South Africa, this resulted in more strategic planning prior to the areas being established. The consequences of establishing these protected areas in the current locations resulted in there being fewer consequences for management today than either Period 1 or 2, with the only consequence being intensive management. The manipulation of the animal populations and vegetation is often mentioned when it comes to management planning as the primary objective of these protected areas is the protection of a specific species, a habitat or for a cultural reason. Over the past few decades there has been a global shift in policy thinking concerning conservation and protected areas, which now emphasises democracy, environmental justice, local involvement and development (Adams & Jeanrenaud, 2008; Rao et al., 2009; Dahlberg et al., 2010), which have ramifications for protected areas managers of these protected areas today. The current main attractions for these protected areas include conservation biodiversity which includes special species, scenery, culture, birding and flora. Owing to these protected areas being owned primarily by government or not for profit organizations, the areas are not as intensively driven by economic survival as there is funding to ensure the longevity of these protected areas.

The majority of manipulations of animal populations and vegetation in Period 2 and 3, include but are not limited to bush clearing and control, alien plant control, supplementary feeding, lick blocks, fire management or vegetation restoration. Only one protected area refers to the manipulation of old agricultural lands with the establishment of grazing lawns as an official conservation project, in an attempt to manipulate the vegetation and its nutritional value for the benefit of large herbivores, and is typically an area established in Period 2.

According to research done by Carruthers (1988) and Hitchcock (2019), conservation approaches in Southern Africa have varied from strict preservationist approaches to community-based conservation. Sandbrook et al. (2019) have shown in their study that there are three independent dimensions of conservation thinking: 1) people-centred conservation – community-based conservation supported by social scientists; 2) science-led ecocentrism – traditional conservation supported by biological scientists and 3) conservation through capitalism – new conservation thinking. There is a lot of speculation about competing goals between the three proponents as to why, what and how to conserve natural areas going forward into the future. The new conservationists argue that the natural capital approach and the use of market-based tools is the way to protect biodiversity, whilst the traditional conservationists argue that we should protect nature for its own sake. The social scientists, or community-based conservationists argue that conservation should be for the benefit of people (Sandbrook et al., 2019). The one area where all three proponents have consensus was that the maintenance of biodiversity and ecosystem processes need to be goals of conservation and that local communities need to benefit from protected areas. The analysis of the selected protected areas showed that with the early establishment of protected areas minimal thought was given to establishing the areas for biodiversity, ecosystems and ignored the interests of the local people in or around the areas, which aligns with Dahlberg et al. (2010). This has changed significantly over the years and the newer protected areas are established on the principles of biodiversity and ecosystems with an emphasis on local communities and their involvement and benefit from protected areas, coupled with the fact that there must be a relationship between conservation, corporations and capitalism.

There is evidence around many protected areas, that the most significant consequences of population growth are the loss and fragmentation of natural habitats through conversion to land uses which support relatively low levels of biodiversity (Ajathi & Krumme, 2002). The world has changed considerably over the past 100 years since the establishment of the first protected areas. Currently, conservation of established or new protected areas has to deal with the rapidly growing human population in the world but especially Africa. For example, the population of Southern Africa has grown from 48 million in 1960 to 212 million in 2020 and Eastern Africa has grown from 68 million in 1960 to 374 million in 2020 (STATS SA, 2020; Trading Economics, 2020; World Bank, 2020). The World Bank believes that over the next 30 years the population will increase to approximately 385 million in Southern Africa and 705 million in Eastern Africa (World Bank, 2020). Since the 1970s, agriculture for food has increased by 300% on earth, with 55% of the oceans being used for fishing (Buitenland, 2019). The increase in the human population leads to more pressure on productive land both for agriculture (the supply of food), land for housing, businesses and bioenergy demands (Anderson-Teixeira et al., 2012). These land uses will be in direct competition with the land currently being used for conservation (Emmott, 2013). The ‘people factor’ in Africa’s current wildlife crisis is by far the most important of all issues involved for protected area conservation (Thomson, 2003; Emmott, 2013).

## Implications for Conservation

Our study suggests that in Southern and Eastern Africa, conservation is on the rebound from poor pre-colonial decisions based on political power, land grabs, uncontrolled hunting, stock farming, agriculture, disease control, mining, a lack of proper conservation knowledge and the disregard for local African knowledge of wild animals. To ensure the long-term survival of protected areas in an era with vastly growing human populations and the ensuing need for food and development, one needs to establish ways of making protected areas on marginal land sustainable for conservation. A further or alternative solution would be to de-gazette land afforded conservation status from prime agricultural land, but this is a far more complex option.

## Supporting information

Supporting Documentation A

## Appendix A

Protected areas in which animal populations or vegetation are actively managed (‘manipulated’). Of the 48 protected areas, we could not obtain information about active management or absence of it for 9 of these areas (i.e., “unestablished” in the Table). The ratio per period indicates whether manipulation was “preferred” (>1) or “avoided” (<1). The three identified periods of establishment are given as P1, P2 and P3

| Manipulation utilised | Of the 48 protected areas identified to be actively/ minimally involved or unestablished with respect to manipulations | The protected areas using manipulations for each period and the relevant percentage | Ratio per period indicating whether manipulation was "preferred" ($>1$) or "avoided" ($<1$) |
| --- | --- | --- | --- |
| Yes | 37 | P1 : 19%<br>P2 : 57%<br>P3 : 24% | P1 : 0.65<br>P2 : 1.14<br>P3 : 1.16 |
| Minimal | 2 | P1 : 50%<br>P3 : 50% | P1 : 1.72<br>P3 : 2.38 |
| Participation in manipulation unestablished | 9 | P1 : 67%<br>P2 : 33% | P1 : 2.30<br>P2 : 0.67 |

## References

Anderson-Teixeira, K. J., Duval, B. D., Long, S. P., & DeLucia, E. H. (2012). Biofuels on the landscape: Is “land sharing” preferable to “land sparing”? Ecological Applications 22(8): 2035–2048. 10.1890/12-0711.1

Carruthers, J. (1992). The Dongola Wild Life Sanctuary: ‘psychological blunder, economic folly and political monstrosity’ or ‘more valuable than rubies and gold’?. African Historical Review Kleio 24: 82–100. 10.1080/00232089285310081

Carruthers, E. J. (2008). “Wilding the farm or farming the wild”? The evolution of scientific game ranching in South Africa from the 1960s to the present. Transactions of the Royal Society of South Africa. Volume 63(2): 160–181. 10.1080/00359190809519220

Child, B. A., Musengezi, J., Parent, G. D., & Child, G. F. T. (2012). The economics and institutional economics of wildlife on private land in Africa. Pastoralism: Research, Policy and Practice, 2:18. http://www.pastoralismjournal.com/content/2/1/18

Child, G., & Riney, T. (1987). Tsetse control hunting in Zimbabwe, 1919–1958. Zambezia XIV: 11–71. PLOS ONE 10.13.71/journal.pone.0081761

Cousins, J. A., Sadler, J. P., & Evans, J. (2008). Exploring the Role of Private Wildlife Ranching as a Conservation Tool in South Africa: Stakeholder Perspectives. Ecology and Society 13 (2): 43. www.ecologyandsociety.org/vol13/iss2/art43

Dahlberg, A., Rohde, R., & Sandell, K. (2010). National Parks and Environmental Justice: Comparing Access Rights and Ideological Legacies in Three Countries. Conservation and Society 8: 209–224. https://www.jstor.org/stable/26393012

Duffy, R. (2014). Waging a war to save biodiversity: the rise of militarized conservation. International Affairs 90: 819–834. 10.1111/1468-2346.12142

Enghoff, M. (1990). Wildlife Conservation, Ecological Strategies and Pastoral Communities. A contribution to the understanding of Parks and People in East Africa. Journal Nomadic Peoples 25-27, 93–107. https://www.jstor.org/stable/43123310

Elton, F. (1873). Journal of an Exploration of the Limpopo River. The Journal of the Royal Geographical Society 42, 1–49. https://www.jstor.org/stable/1798590

Gibson, C. C., & Marks, S. A. (1995). Transforming rural hunters into conservationists: An assessment of community-based wildlife management programs in Africa. World Development 23: 941–957. 10.1016/0305-750X(95)00025-8

Grobler, H. (1996). Dissecting the Kruger myth with blunt instruments: a rebuttal of Jane Carruthers’s view. Southern African Studies 22: 355–472. 10.1080/03057079608708505

Groen, T. A., van Langevelde, F., van de Vijver, C. A. D. M., Govender, N., & Prins, H. H. T. (2008). Soil clay content and fire frequency affecting clustering in trees in South African savannas. Tropical Ecology 24: 269–279. https://www.jstor.org/stable/25172924

Hilbers, J. P., van Langevelde, F., Prins, H. H. T., Grant, C. C., Peel, M., Coughenour, M. B., de Knegt, H. J., Slotow, R., Smit, I., Kiker, G. A., & de Boer, W. F. (2015). Modelling elephant-mediated cascading effects of water point closure. Ecological Applications 25: 402–415. 10.1890/14-0322.1

Hitchcock, R. K. (2001). Hunting is our Heritage: The struggle for hunting and gathering rights among the San of Southern Africa. Senri Ethnological Studies, 59: 139–156. 10.15021/00002791

Hitchcock, R. K. (2019). The Impacts of Conservation and Militarization on Indigenous Peoples. A Southern African San Perspective. Human Nature, 30(1): 217–241. 10.1007/s12110-019-09339-3

Huang, Z. Y. X., van Langevelde, F., Estrada-Pena, A., Suzan, G., & De Boer, W. F. (2016). The diversity-disease relationship: evidence for and criticism of the dilution effect. Parasitology, 143, 1075–1086. Cambridge University Press. 10.1016/S0031182016000536

Kepe, T., Wynberg, R., & Ellis, W. (2005). Land reform and biodiversity conservation in South Africa: complementary or in conflict? International Journal of Biodiversity Science and Management 1: 3–16. 10.1080/17451590509618075

Kirby, C. & E. Kirby. (1931). The Stuart game prerogative. The English Historical Review 46: 239–254. https://www.jstor.org/stable/552949

Klop, E., & Prins, H. H. T. (2008). Diversity and species composition of West African ungulate assemblages: effects of fire, climate and soil. Global Ecology and Biogeography 17: 778–787. 10.1111/j.1466-8238.2008.00416.x

Marston, A. (1996). Aquaculture and the Public Trust Doctrine: Accommodating competing uses of coastal waters in New England. Vermont Law Review 21: 335–374.

Masocha, M., Skidmore, A. K., Poshiwa, X., & Prins, H. H. T. (2011). Frequent burning promotes invasions of alien plants into a mesic African savanna. Biological Invasions 13: 1641–1648. 10.1007/s10530-010-9921-6

Mbaiwa, J. E., & Mbaiwa, O. I. (2006). The effects of veterinary fences on wildlife populations in Okavango Delta, Botswana. International Journal of Wilderness 12: 17–41. http://hdl.handle.net/10311/28

Muir-Leresche, K., & Nelson, R. H. (2000). Private Property Rights to Wildlife: The Southern African Experiment. Competitive Enterprise Institute: 1–31. www.cei.org

Mutwira, R. (1989). Southern Rhodesian Wildlife Policy (1890-1953): A Question of Condoning Game Slaughter? Journal of Southern African Studies 15: 250–262. 10.1080/03057078908708199

Nash, D. J., & Endfield, G. H. (2002). A 19^th^ Century climate chronology for the Kalahari region of central southern Africa derived from missionary correspondence. International Journal of Climatology 22: 821–841. 10.1002/joc.753

Neumann, R. P. (2002). The Post-war Conservation Boom in British Colonial Africa. Environmental History; 7: 22–47. https://www.jstor.org/stable/3985451

Ottichilo, W. K., de Leeuw, J., Skidmore, A. K., Prins, H. H. T., & Said, M. Y. (2000). Population trends of large non-migratory wild herbivores and livestock in the Masai Mara ecosystem, Kenya, between 1977 and 1997. East African Wildlife Society, African Journal of Ecology 38: 202–216. 10.1046/j.1365-2028.2000.00242.x

Ottichilo, W. K., de Leeuw, J., & Prins, H. H. T. (2001). Population trends of resident wildebeest (Connochaetes taurinus hecki) and factors influencing them in the Masai Mara ecosystem, Kenya. Biological Conservation 97: 271–282. 10.1016/S0006-3207(00)00090-2

Plug, I., Scott, K., & Fish, W. (2000). Schoemansdal: faunal remains from selected sites in an historic village. Annals of the Transvaal Museum 37: 125–130. https://hdl.handle.net/10520/AJA00411752_47

Rangarajan, M. (2003). Parks, Politics and History: Conservation Dilemmas in Africa. Conservation and Society 1: 77–98. https://www.jstor.org/stable/26396455

Ripple, W. J., Newsome, T. M., Wolf, C., Dirzo, R., Everatt, K. T., Galetti, M., Hayward, M. W., Kerley, G. I. H., Levi, T., Lindsey, P. A., Macdonald, D. W., Malhi, Y., Painter, L. E., Sandom, C. J., Terborgh, J., & van Valkenburgh, B. (2015). Collapse of the world’s largest herbivores. Science Advances 1: 1–12. DOI: 10.1126/sciadv.1400103

Sandbrook, C., Fisher, J. A., Holmes, G., Luque-Lora, R., & Keane, A. (2019). The global conservation movement is diverse but not divided. Nature Sustainability 2:316–323. 10.1038/s41893-019-0267-5

Scholes, R. J., & Archer, S. R. (1997). Tree–grass interactions in savannas. Annual Review of Ecology and Systematics 28: 517–544. 10.1146/annurev.ecolsys.28.1.517

Spinage, C. A. (1973). A review of ivory exploitation and elephant population trends in Africa. East African Wildlife Journal 11: 281–289. 10.1111/j.1365-2028.1973.tb00093.x

Taylor, R. D., & Martin, R. B. (1987). Effects of veterinary fences in Zimbabwe. Environmental Management 11: 327–334. DOI: 10.1007/BF01867160

Waithaka, J. (2012). Historical facts that shaped Wildlife Conservation in Kenya. The Kenya Wildlife Service in the 21st century: Protecting globally significant areas and resources. The George Wright Forum 29: 21-29. World Journal of Agricultural Research 4: 43–48. https://www.jstor.org/stable/43598971

Waller, R. (1985). Ecology, Migration, and Expansion in East Africa. African Affairs 84: 347–370. 10.1093/oxfordjournals.afraf.a097698

Beard, P. H. (1988). The End of the Game. Chronicle Books, San Francisco, USA.

Cioc, M. (2009). The game of conservation. International treaties to protect the world’s migratory animals. Ohio University Press Books. USA.

Culhane, D. (1998). The pleasure of the crown: Anthropology, law, and First Nations, 120. Talon books. Burnaby, Canada.

Emmott, S. (2013). Ten Billion. Penguin Books. London.

Giles-Vernick, T. (2011). Oral histories: Oral histories as methods and sources. In: E. Perecman & S. R. Curran (eds.). A handbook for social science field research: Essays and bibliographic sources on research design and methods (pp. 85–95). SAGE Publications, London.

Heijnsbergen, V. P. (1997). International legal protection of wild fauna and flora. Landsdale IOS Press. Amsterdam, the Netherlands.

Huntley, B. J., & Walker, B. H. (2012). Ecology of tropical savannas. Springer Science and Business Media: Munich, Germany.

Kesteloot, L. (1992). Myth, Epic, and African History. In V. Y. Mudimbe (ed.), The Surreptitious speech: Présence africaine and the politics of otherness, 1947-1987 (pp. 136). University of Chicago Press, USA.

Prins, H. H. T. (1996). Behaviour and Ecology of the African Buffalo: Social inequality and decision making. Chapman & Hall, London.

Sankaran, M., & Anderson, T. M. (2009). Management and restoration in African Savannas: Interactions and feedbacks. In R. J. Hobbs, & K. N. Suding (eds.). New Models for Ecosystem Dynamics and Restoration (pp. 136–155). Island Press. Washington DC.

Shorrocks, B. & Bates, W. (2015). The Biology of African Savannahs. Oxford University Press, UK.

Stevenson-Hamilton, J. (1938). South African Eden: From Sabi Game Reserve to Kruger National Park. Penguin, 2008. (First Published 1938). Johannesburg.

Strang, D. (1996). Contested sovereignty: the social construction of colonial imperialism. In T. J. Biersteker, & C. Weber (eds.) State sovereignty as social construct (pp. 22–49). Cambridge University Press, UK.

Thomson, R. (2003). A Game Warden’s Report. Magron Publishers. South Africa.

Wagner, R. (1980). Zoutpansberg: the dynamics of a hunting frontier 1848-67. In S. Marks, & A. Atmore (eds.) Economy and Society in Pre-Industrial South Africa (pp. 313–349). Longman, London.

Renwick, A., & Schellhorn, N. (2016). A perspective on land sparing versus land sharing. Learning from agri-environment schemes in Australia: Investing in biodiversity and other ecosystem services on farms (pp. 117–125). Australian National University Press Canberra, Australia.

Worboys, G. L., & Trzyna, T. (2015). ‘Managing Protected Areas’, in G. L. Worboys, M. Lockwood, A. Kothari, S. Feary, & I. Pulsford (eds.). Protected Area Governance and Management (pp. 207–250). ANU Press, Canberra, Australia.

Adams, W. M., & Jeanrenaud, S. J. (2008). Transition to sustainability: Towards a humane and diverse world. Gland: IUCN. https://www.iucn.org/sites/dev/files/import/downloads/transition_to_sustainability_sep_08_en_2.pdf

Department of Environmental Affairs and Tourism. (1997). White Paper on the Conservation and Sustainable Use of South Africa’s Biological Diversity. Government Gazette, 28 July 1997. No. 18163. https://www.environment.gov.za/sites/default/files/legislations/biodiversity_whitepaper_18163_gen1095.pdf

James, A. N., Green, M. J. B., & Paine, J. R. (1999). A Global Review of Protected Area Budgets and Staffing. World Conservation Monitoring Centre. World Conservation Press. Cambridge, U.K. https://www.cbd.int/financial/expenditure/g-spendingglobal-wcmc.pdf

Rao, M., Naro-Maciel, E., & Sterling, E. J. 2009. Protected Areas and Biodiversity Conservation ll: Management and Effectiveness. Network of Conservation Educators and Practitioners (ncep.amnh.org). https://www.researchgate.net/publication/259266417_Protected_Areas_and_Biodiversity_Conservation_II_Management_and_Effectiveness

Watterson, G. G. (1961). Conservation of Nature and Natural Resources in modern African States. IUCN Publications 1. William Clowes and sons, Limited. London. https://www.iucn.org/content/conservation-nature-and-natural-resources-modern-african-states-report-a-symposium

Ajathi, H. M., & Krumme, K. (2002). Ecosystem Based Conservation Strategy for Protected Areas in Savannas. With special reference to East Africa. Thesis. Essen University, Germany. https://nbn-resolving.org/urn:nbn:de:hbz:464-20120803-162654-3

Carruthers, E. J. (1988). Game Protection in the Transvaal 1846 to 1926. Dissertation. University of Cape Town, South Africa. https://open.uct.ac.za/bitstream/handle/11427/23736/Carruthers_Game_protection_in_1988_1.pdf?sequence=1&isAllowed=y

Zulu, N. (2015). An analysis of the post 1980s transition from pastoral to game farming in South Africa: a case study of the Marico district. Dissertation. University of Witwatersrand, South Africa. http://wiredspace.wits.ac.za/handle/10539/19985

Pangeti, G., & Manyanga, M. (n.d.). The Antiquity of Hunting in Southern Africa: The Past to the Present. https://www.academia.edu/2284761/The_Antiquity_of_Hunting_in_southern_Africa_The_Past_to_the_Present. www.5.msu.ac.zw.

Murombedzi, J. C. (2003). Pre-colonial and colonial conservation practices in Southern Africa and their legacy today. Unpublished IUCN manuscript. http://www.iccaconsortium.org

Buitenland. (2019). Krimpende biodiversiteit bedreigt menselijk leven, waarschuwt. https://nos.nl/artikel/2283456-krimpende-biodiversiteit-bedreigt-menselijk-leven-waarschuwt-onderzoek.html

London Gazette. (1889). Charter of the British South African Company. www.rhodesia.me.uk/charter

Oxford Dictionary. (2017, January 5).www.en.oxforddictionaries.com/definition/res_nullius

Paterson, A. R. (2009). *Legal Framework for Protected Areas: South Africa*. www.iucn.org/downloads.South_Africa

STATS SA. (2020). Statistics South Africa. The South Africa I know, the home I understand. http://www.statssa.gov.za/?page_id=593

Trading Economics. (2020). South Africa – Economic Indicators. https://tradingeconomics.com/south-africa/indicators

Vidal, J. (2011). Africa’s declining wildlife. The Guardian. www.theguardian.com/environment/2011/jun/20/africa-declining-wildlife_12

World Bank. (2020). Population Estimates and Projections. https://datacatalog.worldbank.org/dataset/population-estimates-and-projections

